# Colony phase structure favors permanent worker evolution in juvenile social systems

**DOI:** 10.64898/2026.04.24.720620

**Authors:** Nobuaki Mizumoto

## Abstract

Permanent workers in social insects, who forgo reproduction to help others, are a defining feature of superorganisms and a major evolutionary transition. Most theories assume that helping in the natal nests is inherently mutually exclusive from dispersal to found a new colony. This assumption holds for many adult societies whose workers cannot molt and change castes (e.g., Hymenoptera), but not for juvenile societies whose workers may differentiate and disperse after a period of helping (e.g., termites). The evolutionary advantage of permanent workers remains unknown in such juvenile societies with developmental flexibility. Here we develop a demographic model in which individuals can disperse either before or after a period of helping, capturing reversible helpers and permanent workers in termites. We found that permanent workers are favored when colony growth is divided into distinct ergonomic and reproductive phases, even if their performance is the same as that of reversible helpers. Without this phase separation, there is no selective advantage for workers to lose dispersal for colony foundation. After the phase separation, on the other hand, permanent workers increase continuous demographic growth and evolve through kin selection. By mapping diverse termite social systems onto a continuous landscape of ontogeny, the model traces a pathway linking different social forms. These results generalize social evolution theory beyond adult-worker systems by providing a demographic mechanism that favors the loss of the reproductive option.

## Introduction

The evolution of the reproductive division of labor is a major transition in social evolution (1–3). Permanent workers have evolved independently in several lineages, including social Hymenoptera, ambrosia beetles, and termites, through diverse evolutionary pathways (4–8). Most theoretical models assume that becoming a worker or a reproductive is a mutually exclusive option (9–13). This assumption reflects adult-worker systems, such as Hymenoptera and beetles, where workers cannot further differentiate (14–16), with some exceptions (17, 18). However, this framework does not apply to juvenile worker systems, such as termites. Termite workers are immature and may retain the potential to disperse later due to hemimetabolous development (19–21). How permanent workers evolve from such flexible developmental systems remains unresolved.

All termite societies include workers, and their reproductive potential depends on the species (22–24). Observations of wood-feeding cockroaches suggest that the ancestor was a subsocial family with biparental care, without workers (Fig. 1). Termite workers are classically categorized as pseudergate or true workers (Fig. 1). In pseudergate systems, workers can molt into alates to disperse and found new colonies. In contrast, true workers are analogous to hymenopteran workers, as they irreversibly diverge from reproductive development. Thus, pseudergates contribute to both colony growth and future reproduction, while true workers do not. This raises a question: what is the selective advantage of losing the option to disperse after the period of helping?

**Figure 1.**
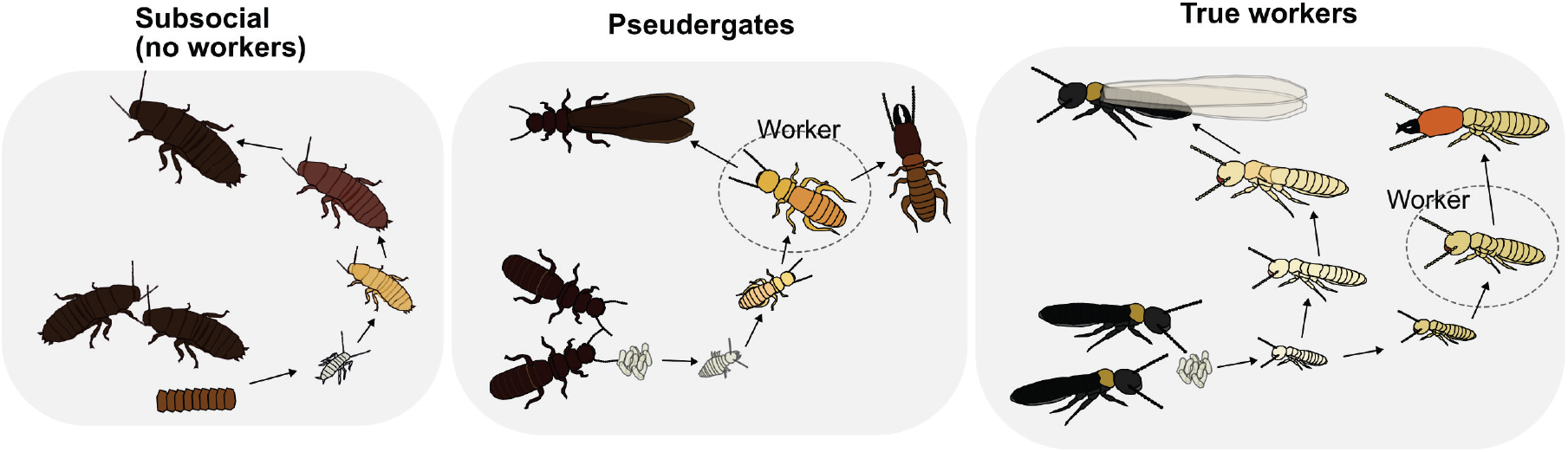
Classic worker types in termites. In the pseudergate society, workers can differentiate alates to reproduce colonies. On the other hand, in the true worker society, workers irreversibly diverged from the alate line.

Despite repeated calls, theoretical models for termite worker evolution remain limited (19, 20, 25). Because true workers express developmental canalization during immature development, true worker evolution must be a result of the long-term continued selection of helping without dispersal under the genetic monogamy (26). Then, under which conditions is such behavior to permanently remain in the nest favored in the first place? Higashi et al. 1991 predicted that foraging is a decisive factor in the evolution of permanent workers by forming long-lasting nests (27). However, there are two imitations in this model. First, it assumes that permanent workers are inherently much more efficient helpers than facultative helpers, which is unlikely at early evolutionary stages when phenotypic differences should be minimal (24, 28, 29). Second, although many previous studies assumed that foraging automatically led to true worker evolution (19, 20, 27, 30–32), these are not necessarily associated with each other. At least six genera (four families) have pseudergates that forage outside the nest (33–40).

These limitations suggest that factors beyond helper efficiency and foraging ecology are required to explain the evolution of permanent workers. Colony demography may play a critical role in distinguishing the function of pseudergates and true worker societies. In social insects with permanent workers, colony growth is divided into an ergonomic phase focused on worker production and a reproductive phase focused on alate production (41). As such, in true worker termites, the alate production line is independent of the worker production line. In contrast, in pseudergate societies, the workload is directly converted into alates. This difference implies that the organization of colony growth may determine whether the behavior to permanently remain in the nest is favored. Because previous life history theories of social insects assume colonies with permanent workers (41, 42), it remains unclear how the developmental schedule itself can facilitate permanent worker evolution.

Here we develop a demographic framework that treats worker fate as a continuous trait, allowing individuals to disperse either before or after a helping period. By modeling developmental pathways as quantitative traits, we capture the full range of termite social systems (Fig. 2). We show that permanent workers are favored only when colony growth is divided into ergonomic and reproductive phases. Before this phase separation, selection favors delayed dispersal but not permanent workers. Our results identify that colony growth structure as a mechanism linking reversible and irreversible social systems.

**Figure 2.**
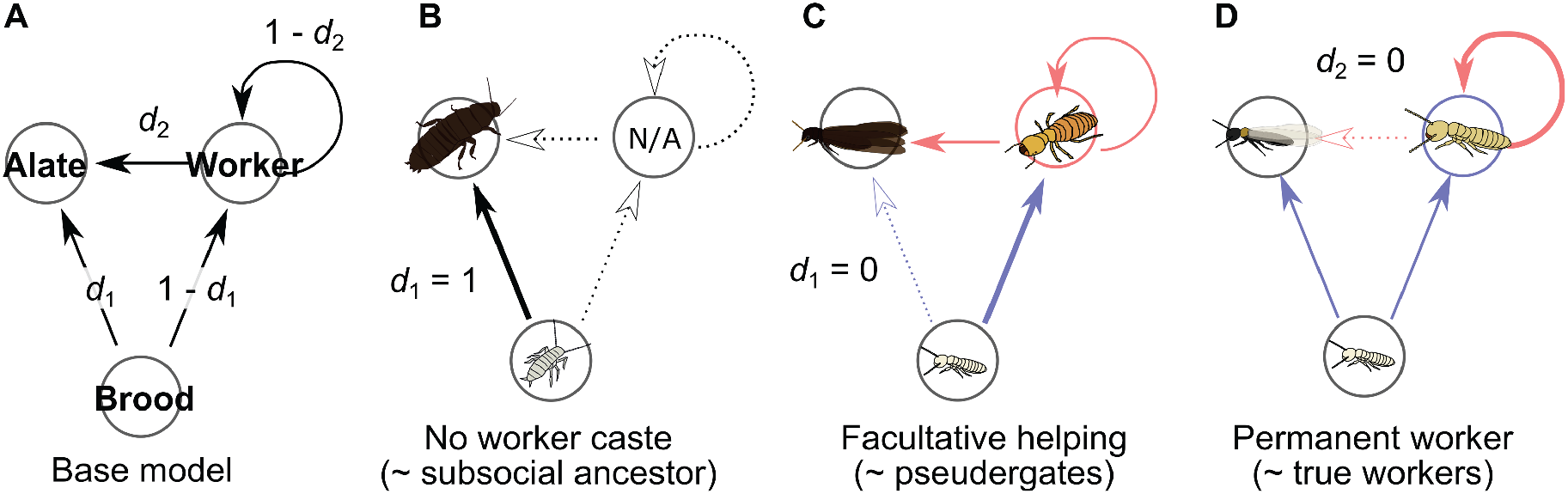
Model framework. (A) Our framework considers a simplified developmental pathway among brood, workers, and alates. Brood can differentiate either workers (the probability of 1–*d*_1_) or alates (*d*_1_), while workers may remain workers (1–*d*_2_) or differentiate into alates (*d*_2_). (B) Subsocial species without worker castes can be described by *d*_1_ = 1, where all brood invests into maturation without contributing to further brood production. (C) In pseudergate societies, broods usually differentiate into workers, and then workers can become alates (*d*_1_ = 0 and *d*_2_ > 0). (D) In true worker societies, broods either differentiate into workers or alates, but workers do not differentiate into alates (*d*_1_ > 0 and *d*_2_ = 0).

## Results

### A continuous framework for termite worker status

Our framework treats discrete social systems (subsocial, pseudergates, and true workers) as quantitatively differential expressions of the shared transition probabilities (Fig. 2A). We consider simply three developmental stages: brood (predevelopmental), workers (contribute to the brood production by helping), and alates (disperse to start new colonies). Brood and workers can become alates to disperse with probabilities *d*_1_ and *d*_2_, respectively, or otherwise remain in the nest to help sibling production. In subsocial species without worker castes, all broods directly molt into dispersers (*d*_1_ = 1, Fig. 2B). In facultative helping society, broods usually molt into workers and then workers become alates, similar to pseudergate termite species (*d*_1_ = 0 and *d*_2_ > 0, Fig. 2C). In permanent worker societies, broods either differentiate into workers or alates, but workers do not differentiate into alates, similar to true worker termite species (*d*_1_ > 0 and *d*_2_ = 0, Fig. 2D). This formulation allows us to treat different social systems within a single continuous framework of two dispersal parameters, *d*_1_ and *d*_2_, enabling direct comparison of their evolutionary dynamics.

### Analytical model: true workers are favored only with the colony demographic schedule

Without a distinct ergonomic stage, a permanent worker strategy was never favored over facultative helping (Fig. 3A-C; Fig. S1). Even when the worker contribution is extremely high (*f* > 1), where workers contribute to producing more than one worker per season, facultative helping is better than permanent workers (Fig. S3), since there is no reason for workers to lose their reproductive potential. Even with foraging behavior, characterized by long colony life (large *T*_max_) and low worker survival (small *s*), facultative helping societies remain advantageous (Fig. S2). Thus, foraging alone is not enough for the evolution of permanent workers, consistent with empirical observations (33–39).

**Figure 3.**
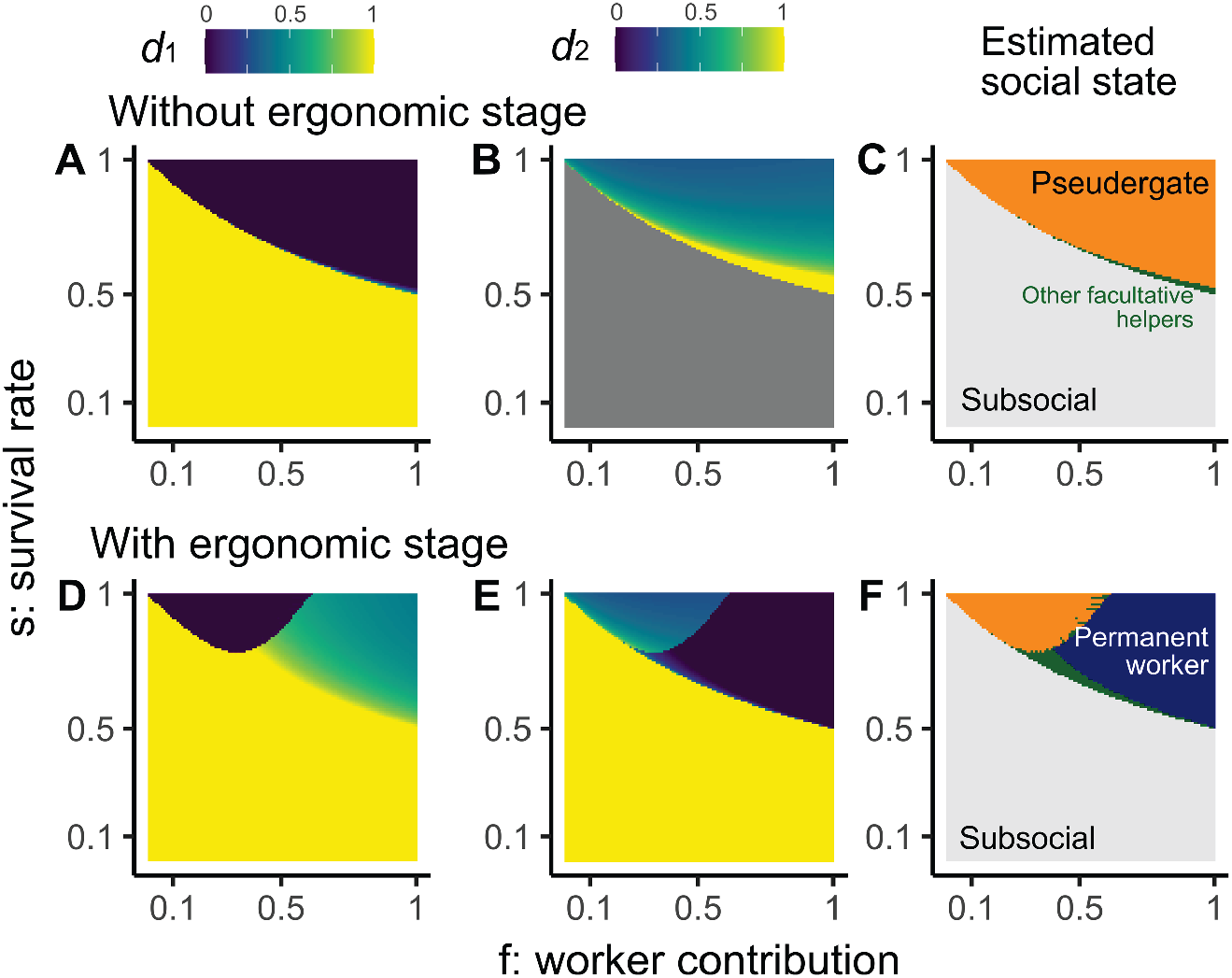
Analytical solutions for the most advantageous social strategy across different parameters, *s* and *f*. The social states (C, F) are estimated from *d*_1_ (A, D) and *d*_2_ (B, E), with subsocial being *d*_1_ = 1, pseudergate being *d*_1_ = 0 and *d*_2_ > 0, and permanent worker being *d*_1_ > 0 and *d*_2_ = 0. Others are regarded as facultative helter types, as *d*_2_ > 0.

In contrast, when colony growth is divided into ergonomic and reproductive phases, permanent workers can become advantageous (Fig. 3D-F). During the ergonomic stage, workers accumulate in the colony in both systems. Once the colony switches to the reproductive phase, pseudergate-colonies with facultative helpters produce alates from workers, while true worker colonies with permanent workers reproduce alates from broods. The former can produce large amounts of alates in short periods at the cost of reducing the working force. The latter produces alates more slowly at the beginning but can continue to increase in the long term, as the workload will not be reduced. In this scenario, permanent workers are more advantageous than facultative helpers when worker contribution is larger (Fig. 3F). Interestingly, with the lower worker contribution (e.g., *f* = 0.5), permanent workers are favored when the survival rate is lower, which should reflect foraging behaviors. This is because, in pseudergate societies, the lower survival rates of workers directly translate into lower survival rates of alates. Thus, our results are consistent with the classic prediction of termite worker evolution: the association between foragers and true workers (19, 27, 30), but also highlight that this association is possible after the evolution of a clear reproductive schedule.

### Evolutionary simulations: true workers evolved from a pseudergate society

Evolutionary simulation confirmed the analytical prediction under the explicit monogamous mating conditions. Without ergonomic stages, facultative helper society with the pseudergates (*d*_1_ = 0 and *d*_2_ > 0) evolved from a subsocial ancestor, and permanent workers did not evolve, even with high worker contributions (Fig. 4AB). In contrast, by introducing a colony reproductive schedule of the ergonomic stage, we observed the evolution of permanent workers (Fig. 4D). When the worker contribution was small, evolutionary simulations end up with the state of pseudergate society (e.g., *f* = 0.25; Fig 4C), while with high worker contributions, permanent worker society evolved from the initial state of subsocial colonies (*f* = 1; Fig. 4D).

**Figure 4.**
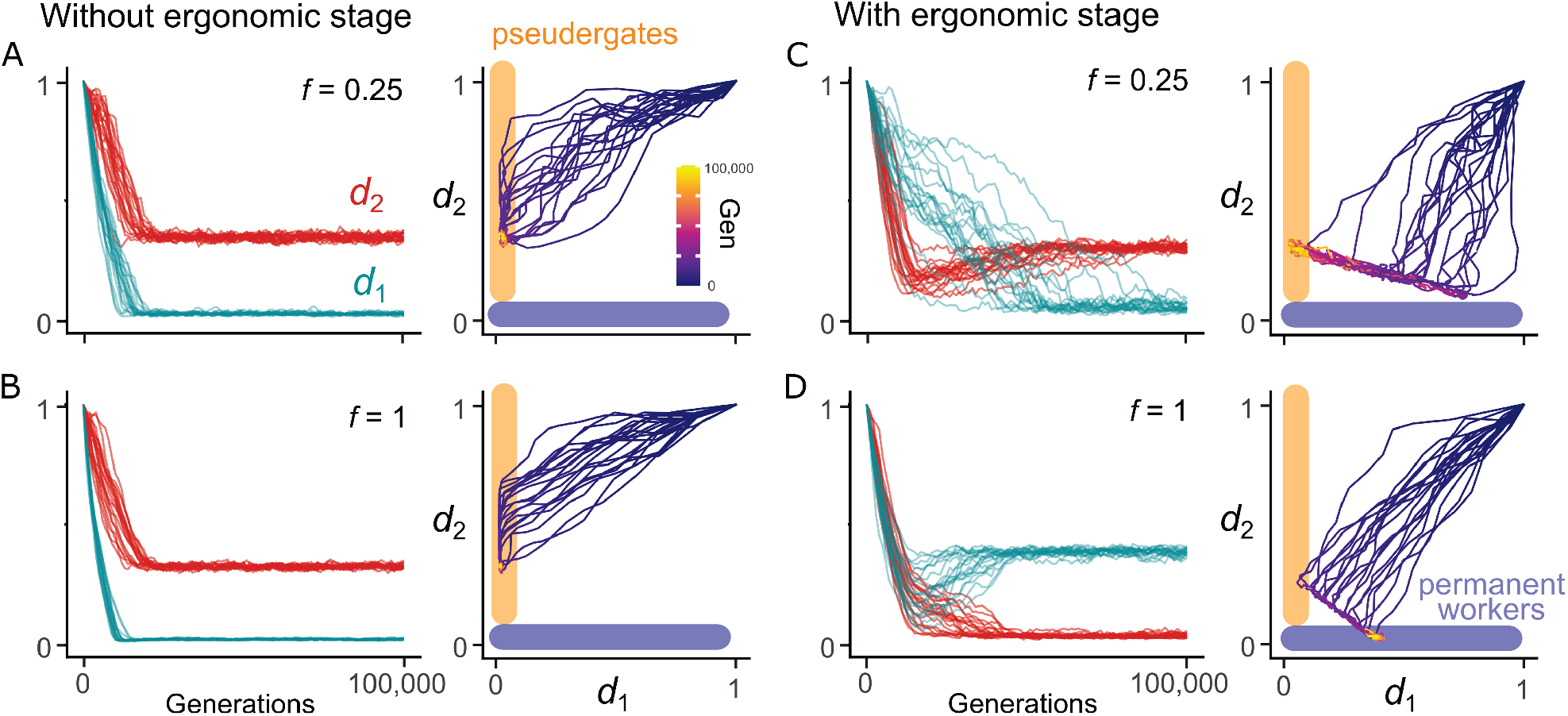
Evolution of true workers with a colony demographic schedule. The lines indicate the 20 evolutionary trajectories of mean trait values (*s* = 0.95, *T*_max_ = 10, *N* = 5,000, σ_mutation_ = 0.002). The evolution of *d*_1_ and *d*_2_ was mapped into the landscape of termite social systems.

Importantly, the evolutionary trajectories reveal a gradual transition from subsocial to facultative helping (even through pseudergates) to permanent workers. Without the ergonomic phase, all evolutionary trajectories directly moved from the subsocial ancestor towards pseudergate society (Fig. 4 AB). On the other hand, with the ergonomic stage, the evolutionary trajectories were more complex. The population first quickly moved from the ancestral subsocial state to the attractor line with high fitness in the parameter landscape (Fig. 4CD, Fig. S4). Then, it moved along the line toward either the optimal pseudergate or permanent worker societies. Several evolutionary simulations clearly followed the path from a subsocial ancestor to a pseudergate society, then moved toward a permanent worker society (Fig. 4D).

### Evolution of phase separation

Finally, we directly investigated the advantage of an ergonomic stage by comparing societies with and without the separation of the ergonomic and reproductive phases. Our analytical solution shows that the pseudergate society can become the most advantageous strategy either with or without an ergonomic stage (Fig. 5A). However, the permanent worker strategy can become the most advantageous strategy only when there is a separation of ergonomic and reproductive phases (Fig. 5A). Evolutionary simulations of phase separation further showed the relationship between ergonomic phase and social systems. Without phase separation, pseudergate society is an evolutionary stable state across parameters (left side of dashed line in Fig. 5B-D). On the other hand, different dynamics were observed after the mutation of the phase separation was introduced, including the evolution of phase separation without changing social systems (Fig. 5B), the evolution of phase separation followed by the evolution of permanent workers (Fig. 5C), and remaining in the pseudergate society without phase separation (Fig. 5D). Thus, phase separation of ergnomic and reproductive stages can evolve either along with or independent from the evolution of permanent workers.

**Figure 5.**
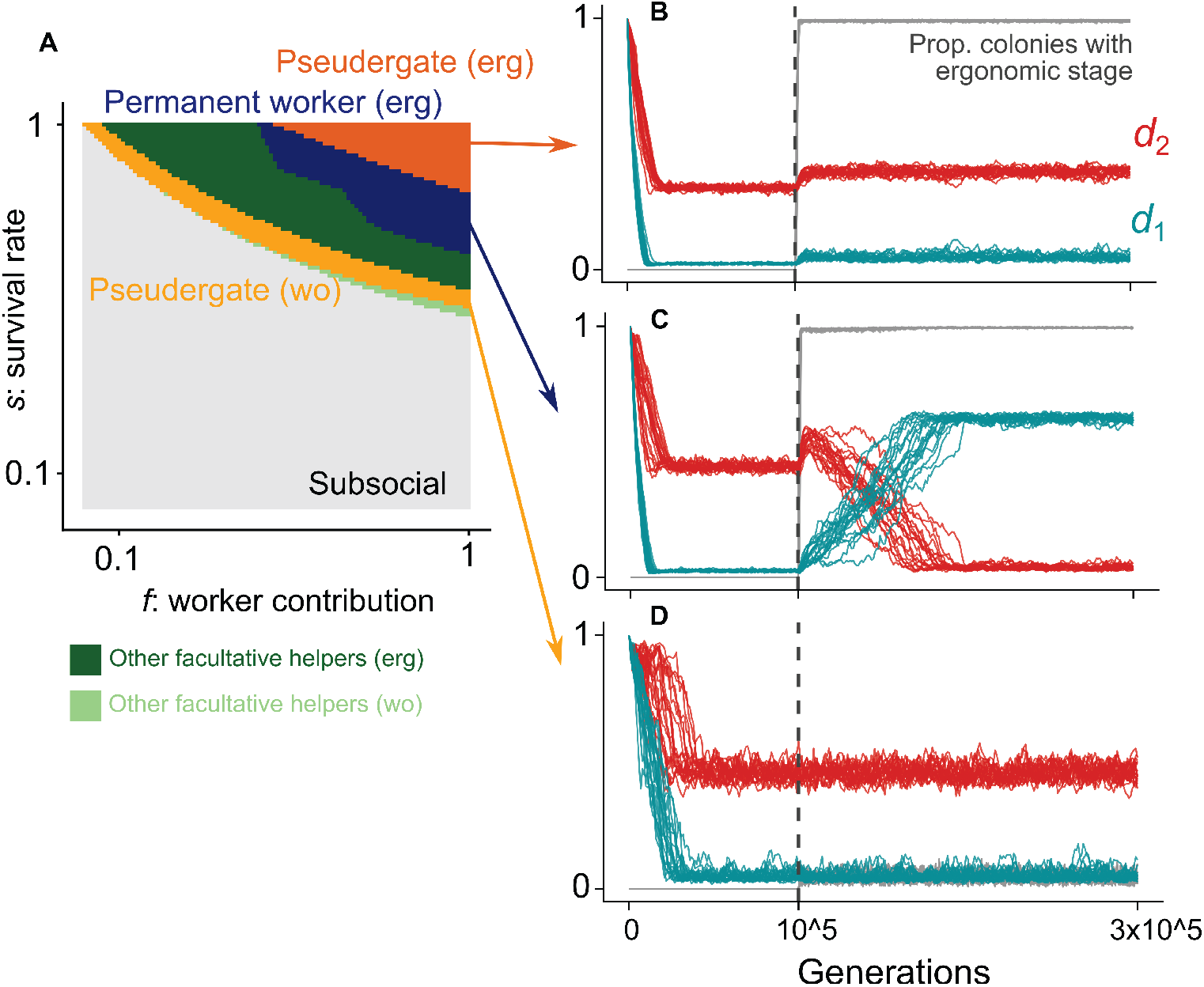
The evolution of phase separation. (A) Analytical solutions for the most advantageous social strategy (described by *d*_1_, *d*_2_, and phase separations). erg indicates the condition with an ergonomic stage, while wo indicates that without an ergonomic stage. (B-D) Evolutionary simulations of the phase separation and social systems. The simulations start from the condition without ergonomic stages. After 10_5_ generations, the phase separation was allowed to evolve (dashed vertical lines). The lines indicate the 20 evolutionary trajectories of mean trait values (B: *s* = 0.95 *f* = 1, C: *s* = 0.75 *f* = 1, D: *s* = 0.95 *f* = 0.1; *T*_max_ = 10, *N* = 5,000, σ_mutation_ = 0.002).

## Discussion

Our results show that permanent workers can be evolutionarily favored without any intrinsic advantage over facultative helpers. Instead, selection can favor permanent workers when colony growth is divided into distinct ergonomic and reproductive phases by optimizing colony demography. This challenges the common assumption that obligate workers must be more efficient than facultative helpers (27). The efficiency of obligate workers should have evolved after the specialization (43, 44), and thus, the phenotypic differences between the two forms should be minimal at the early stage of social evolution. Under such incipient conditions, we show that phase-structured colony growth alone is sufficient for permanent worker evolution. Once some workers remain permanently in the nest, subsequent selection can further canalize the development and differentiation of working individuals in the nest (26), increasing true worker efficiency to the modern termites.

By modeling workers as a continuous trait with two dispersal parameters, our framework demonstrates smooth evolutionary transitions between subsocial systems, pseudergates, and true workers. This perspective helps resolve a long-standing inconsistency in the ancestral state of termite workers. Many consider that pseudergates represent the ancestral state, as both woodroaches and pseudergates have one developmental line (26, 30, 45) (Fig. 1). Others claim the conclusion depends on how developmental pathways are coded; differentiation between winged-wingless dimorphism, similar to alates and workers, could have existed in the ancestor (46–48). Or, the most conservative approach, treating these three as distinct traits, highlights the inconclusiveness of the ancestral state (49). This inconsistency arises from treating worker states as coarse categories, rather than quantitative traits (48). In reality, species differ in the number of molts, developmental flexibility, and pathways to become alates (23, 48, 50). Our approach of mapping social evolution onto a continuous trait landscape, rather than discrete steps, is more operationally suitable to capture the diversity of termite social organization and aligns with similar shifts toward quantitative frameworks in studies of social Hymenoptera (8, 51, 52).

Our model predicts that true worker societies should exhibit a strict ergonomic-reproductive separation, whereas pseudergate societies should show greater diversity in phase separation. This prediction is consistent with empirical observations of alate production timing across termites. In species with true workers, alate production only starts after colonies reach large sizes (on the order of 10^3^-10^6^ termites) (53–55). Even in species with accelerated life history, colonies pass through a distinct ergonomic-reproduction phase (e.g., only 3 years of colony life in *Silvestritermes minutus* (56)). On the other hand, colonies with pseudergates also often undergo ergonomic phases, but the clarity of the distinction largely varies. In one-piece nesting termites, alate production starts progressively at an early stage (57–59) and is often triggered by environmental cues, such as resource depletion (60). On the other hand, foraging pseudergate species have clearer ergonomic and reproductive separation, without nymphal differentiation until they reach a certain colony size (40). Notably, our model detected two distinct selection regimes in pseudergates, with and without the ergonomic stage (Fig. 5), which may reflect differences in foraging strategy observed in pseudergate species.

Note that the permanent worker strategy in our model (*d*_2_ = 0) represents the behavioral strategy for workers to remain in the nest for their entire life, without morphological differentiation. In this sense, permanent workers in our model are different from true workers in the modern termites, which have experienced different degrees of morphological differentiation, depending on lineages (48). Thus, our model is relevant for considering the early stages of worker evolution without morphological differentiation. We found that the benefit of specialization arises from colony growth dynamics, without the intrinsic high efficiency of permanent workers. This suggests the presence of behavioral castes even in early termites or pseudergate societies. Modern termites also show diversity with mixed developmental potentials, e.g., some have both helpers with and without the potential to become alates (39), while some species have behaviorally distinct castes within a single ontogenic process (61). Our framework is flexible to accommodate such variation beyond coarse-graining worker systems, promising future expandability to incorporate more detailed developmental pathways.

In conclusion, we provide a general framework for the evolution of permanent workers in systems with reversible helping by explicitly modeling flexible development. The theories of termite social evolution have been based on verbal models (24, 32, 62, 63), often lacking quantitative tests. Our framework offers a quantitative basis for studying termite social evolution, which can be easily expanded to address additional factors, such as non-genetic monogamy (64), soldier castes (65), worker polyphenism (23, 50), and nest inheritance (24, 66). By linking colony demography with social evolutionary outcomes, our study extends worker evolution theory to juvenile systems and integrates developmental, ecological, and molecular perspectives of termite social organizations.

## Methods

### Basic model

Inspired by the matrix population model framework (67), we used a projection matrix to track changes in colony composition and alate production over time. Each colony comprises the broods, workers, and alates, where reproductives (king and queen) produce broods based on worker contribution. Broods can differentiate into workers or alates, while workers may remain workers or differentiate into alates, based on the dispersal rate of *d*_1_ and *d*_2_ (Fig. 2). Given that the within-colony population at generation *t* is **n**_*t*_ = (*brood*(*t*), worker(*t*), *alates*(*t*))^T^, the population is updated as **n**_*t*+1_ = **A**(*s, f, d*_1_, *d*_2_)**n**_*t*_, expanded as

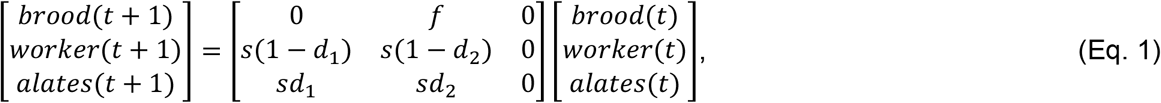

where *s* is the survival rate within a colony, and *f* is the worker contribution towards brood production by helping parents and siblings. The population is updated until it reaches the colony life-span, *T*_max_. This framework enables tracking the temporal development of demography for each colony, including the colony size and alate production. Note that we do not consider the nest inheritance in this study (24). The king and queen live until the end of the colony lifespan.

### Analytical solutions

We tested the advantage of each social structure, described by the combination of *d*_1_ and *d*_2_, across a variety of ecological conditions, represented by combinations of parameters, *s, f*, and *T*_max_ (Table S1). Given that **n**_**0**_ is the initial population and the colony life span is *T*_max_, the colony composition at *T*_max_ is 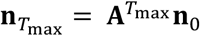, and the cumulative number of produced alates is 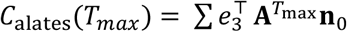, where *e*_3_ = (0,0,1). We consider two initial conditions, one is without a colony developmental schedule, **n**_**0**_ = **n**_**0wo**_ = (10, 0, 0), where the colony starts from only broods; the other is with a colony developmental schedule, **n**_**0**_ = **n**_**0w**_ = (5, 10, 0), where colony went through production of first workers (ergonomic stage) and then start producing alates (reproductive stage). We obtained the cumulative number of produced alates as a measure of the colony performance with a social structure. Note that this projection matrix is reducible and nonergodic (67, 69), different from common matrix population models.

### Evolutionary simulations

We develop an evolutionary simulation by explicitly modeling the monogamous mating system of diploid organisms. Colonies can persist for one or several years based on the colony member dynamics. For example, a colony founded by a monogamous pair of a female (genotype AB) and a male (CD) produces offspring with four possible genotypes: AC, AD, BC, and BD. Broods with the AC genotype can develop only into workers or alates with the AC genotype, but workers contribute equally to the production of all brood types; thus, lower relatedness poses a challenge for worker evolution. In this system, the matrix population model in Eq. 1 can be expanded as follows:

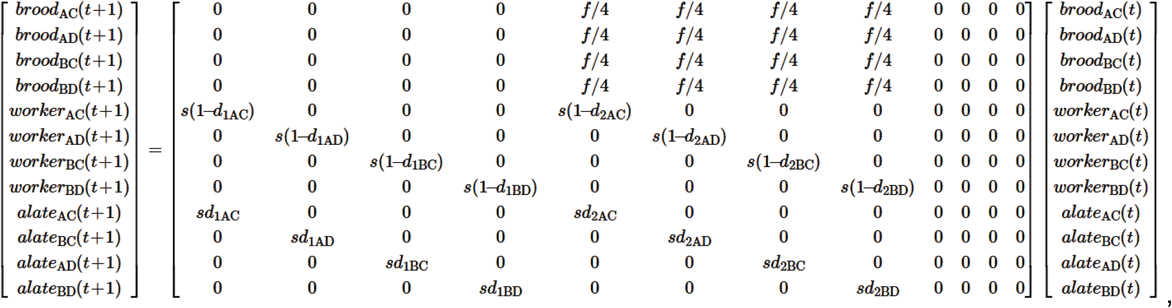

where each genotype has its own unique dispersal probability (e.g., *d*_1AC_, *d*_1AD_, *d*_1BC_, and *d*_1BD_). Lowering dispersal probability is considered an altruistic behavior, as individuals remain in the nest, incur the cost of dying before dispersing, and contribute to the production of siblings. The relatedness between individuals is described by considering the genotype of all individuals (9).

This framework is consistent with the models developed in social evolution in bees, which explicitly modeled the genetic structure for relatedness and used dispersal rates for worker castes (9, 12, 13).

Each simulation starts with *N* identical subsocial nests (*d*_1_ = *d*_2_ = 1). Each colony in the population has its own colony development dynamics based on the genotype combinations of the colony founders. Colonies survive until they reach *T*_max_ or their colony size becomes smaller than 1. All alates in the colony dispersed to found new colonies at available patches within *N*, and alates that failed in colony foundation were dead and did not exist the next year. Both females and males carry two sets of two genes (*d*_1_ and *d*_2_). Mutations occur per locus by adding a normal distribution error with mean 0 and s.d. σ_mutation_. Genes are expressed as the mean value of two alleles. If *d*_1_ and *d*_2_ became less than 0 or larger than 1, they were clamped as 0 or 1, respectively. We tested the evolutionary consequences of four situations: without an ergonomic stage, with low worker contribution *f* and high *f*, and with an ergonomic stage, with low *f* and high *f*. Without an ergonomic stage, each colony starts from 100 broods (25 for each genotype), while with an ergonomic stage, each colony starts from 100 broods (25 for each genotype) and 200 workers (50 for each genotype). We use 0.25 for low *f*, while 1 for high *f*. Other biological parameters, *s* and *T*_max_, were fixed as 0.95 and 10, respectively. We used the population size *N* = 5,000, mutation rate σ_mutation_. = 0.002, and a generation time of 100,000 years (20*N*), which is long enough to achieve an evolutionary plateau in this population size and mutation rates. We performed a sensitivity analysis of these simulation-related parameters, which did not change the conclusion (Fig. S5-6). We ran the simulation for each parameter combination 20 times.

### Evolution of phase separation

We further compare the colony productivity between conditions with and without phase separation. Using the above analytical model without the ergonomic phase (**n**_**0wo**_), we examine the condition where colonies produce alates only after reaching a certain colony size (*n*_thresh_). Before this threshold, both *d*_1_ and *d*_2_ were set to 0, resulting in ergonomic growth. Then, we classified the most productive social structures based on *d*_1_, *d*_2_, and *n*_thresh_. The values of *n*_thresh_ were either 0 (same as the above model without ergonomic phase) or 15 (corresponding to the condition with ergonomic phase in the above model).

Then, we performed evolutionary simulations. In addition to the simulation above, we added another locus that codes for ergonomic growth (0/1); when this gene is expressed, individuals do not become alates (*d*_1_ = *d*_2_ = 0) until the colony reaches the threshold colony size (*n*_thresh_ = 150, corresponding to the condition of analytical solution). We treated this as a recessive trait, with expression only when both alleles are 1. Thus, depending on the parent genotypes, some workers exhibit ergonomic state-like behaviors, while others do not. The evolutionary simulations were performed for 300,000 generations, where the mutation at the ergonomic growth locus became available after 100,000 generations. The mutation occurs in 0.1 %. We used *s* = 0.75, *f* = 1.0; *s* = 0.95, *f* = 1.0; and *s* = 0.95, *f* = 0.1, to represent different evolutionary scenarios (Fig. 5), while other parameters, *T*_max_, = 10, population size *N* = 5,000, and the mutation rate σ_mutation_. = 0.002.

All computational works were performed using R version 4.5.2 (70). The major parts of the evolutionary simulation were constructed with C++, with the *Rcpp* v1.0.14 package integration (71).

## Ackowledgements

We would like to thank Dr. Koos Boomsma and Elijah Carroll for their helpful comments on the earlier version of the draft, and the Auburn University Social Insects Shared meeting for its helpful feedback on the early results.

## Statements & Declaration

### Data Availability

The codes generated during the current study are available on GitHub (https://github.com/nobuaki-mzmt/true-worker-evolution). The accepted version will be deposited in Zenodo to obtain a DOI.

### Ethics approval

This study did not require ethical approval as no new data were collected from human or animal subjects.

## References

1. J. J. Boomsma, R. Gawne, Superorganismality and caste differentiation as points of no return: how the major evolutionary transitions were lost in translation. Biological Reviews 93, 28–54 (2018).

2. S. A. West, R. M. Fisher, A. Gardner, E. T. Kiers, Major evolutionary transitions in individuality. Proceedings of the National Academy of Sciences of the United States of America 112, 10112–10119 (2015).

3. G. A. Cooper, S. A. West, Division of labour and the evolution of extreme specialization. Nature Ecology & Evolution 2, 1161–1167 (2018).

4. R. Gadagkar, Why the Definition of Eusociality Is Not Helpful to Understand Its Evolution and What Should We Do about It. Oikos 70, 485–488 (1994).

5. K. Tsuji, Sterility for life: applying the concept of eusociality. Animal Behaviour 44, 572–573 (1992).

6. H. Burda, R. L. Honeycutt, S. Begall, O. Locker-Grütjen, A. Scharff, Are naked and common mole-rats eusocial and if so, why? Behavioral Ecology and Sociobiology 47, 293–303 (2000).

7. B. J. Crespi, D. Yanega, The definition of eusociality. Behavioral Ecology 6, 109–115 (1995).

8. T. A. Linksvayer, B. R. Johnson, Re-thinking the social ladder approach for elucidating the evolution and molecular basis of insect societies. Current Opinion in Insect Science 34, 123–129 (2019).

9. E. Rees-Baylis, I. Pen, J. J. Kreider, Maternal manipulation of offspring size can trigger the evolution of eusociality in promiscuous species. Proceedings of the National Academy of Sciences of the United States of America 121, e2402179121 (2024).

10. L. Fromhage, H. Kokko, Monogamy and haplodiploidy act in synergy to promote the evolution of eusociality. Nature Communications 2, 397 (2011).

11. P. Nonacs, Monogamy and high relatedness do not preferentially favor the evolution of cooperation. BMC Evolutionary Biology 11, 58 (2011).

12. A. E. Quiñones, I. Pen, A unified model of Hymenopteran preadaptations that trigger the evolutionary transition to eusociality. Nature Communications 8, 15920 (2017).

13. D. M. Ruttenberg, S. A. Levin, N. S. Wingreen, S. D. Kocher, Variation in season length and development time is sufficient to drive the emergence and coexistence of social and solitary behavioural strategies. Proceedings of the Royal Society B: Biological Sciences 291, 20241221 (2024).

14. D. R. Rubenstein, P. Abbot, Eds., Comparative Social Evolution (Cambridge University Press, 2017).

15. E. O. Wilson, Sociobiology: The New Synthesis, Twenty-Fifth Anniversary Edition (Harvard University Press, 2000).

16. S. M. Smith, D. S. Kent, J. J. Boomsma, A. J. Stow, Monogamous sperm storage and permanent worker sterility in a long-lived ambrosia beetle. Nature Ecology & Evolution 2, 1009–1018 (2018).

17. R. Gadagkar, Demographic predisposition to the evolution of eusociality: a hierarchy of models. Proceedings of the National Academy of Sciences 88, 10993–10997 (1991).

18. M. Zhuang, et al., Unexpected worker mating and colony-founding in a superorganism. Nature Communications 14, 5499 (2023).

19. J. Korb, B. Thorne, “Sociality in Termites” in Comparative Social Evolution, Cambridge University Press, D. R. Rubenstein, P. Abbot, Eds. (Cambridge University Press, 2017), pp. 124–153.

20. Y. Roisin, J. Korb, “Social Organisation and the Status of Workers in Termites” in Biology of Termites: A Modern Synthesis, (Springer Netherlands, 2010), pp. 133–164.

21. Y. Roisin, “Diversity and Evolution of Caste Patterns” in Termites: Evolution, Sociality, Symbioses, Ecology, T. Abe, D. E. Bignell, M. Higashi, Eds. (Springer Netherlands, 2000), pp. 95–119.

22. C. Noirot, J. M. Pasteels, Ontogenetic development and evolution of the worker caste in termites. Experientia 43, 851–860 (1987).

23. L. Revely, S. Sumner, P. Eggleton, The plasticity and developmental potential of termites. Frontiers in Ecology and Evolution 9, 1–13 (2021).

24. Y. Roisin, Intragroup Conflicts and the Evolution of Sterile Castes in Termites. The American Naturalist 143, 751–765 (1994).

25. C. A. Nalepa, “Altricial development in wood-feeding cockroaches: the key antecedent of termite eusociality” in Biology of Termites: A Modern Synthesis, D. E. Bignell, Y. Roisin, N. Lo, Eds. (Springer, 2011), pp. 69–96.

26. Y. Cui, et al., Nutritional specialization and social evolution in woodroaches and termites. Science eadt2178 (2026). 10.1126/science.adt2178.

27. M. Higashi, N. Yamamura, T. Abe, T. P. Burns, Why don’t all termite species have a sterile worker caste ? Proceedings of the Royal Society B: Biological Sciences 246, 25–29 (1991).

28. D. Parmentier, Y. Roisin, Caste morphology and development in Termitogeton nr. planus (Insecta, Isoptera, Rhinotermitidae). Journal of Morphology 255, 69–79 (2003).

29. K. Matsuura, et al., Genomic imprinting drives eusociality. [Preprint] (2021). Available at: https://www.biorxiv.org/content/10.1101/2021.04.28.441822v1 [Accessed 21 March 2025].

30. D. J. G. Inward, A. P. Vogler, P. Eggleton, A comprehensive phylogenetic analysis of termites (Isoptera) illuminates key aspects of their evolutionary biology. Molecular Phylogenetics and Evolution 44, 953–967 (2007).

31. T. Abe, “Evolution of life types in termites” in Evolution and Coadaptation in Biotic Communities, S. Kawano, J. Connell, T. Hidaka, Eds. (University of Tokyo Press, 1987), pp. 125–148.

32. J. Korb, “The ecology of social evolution in termites” in Ecology of Social Evolution, J. Korb, J. Heinze, Eds. (Springer Berlin Heidelberg, 2008), pp. 151–174.

33. N. Mizumoto, T. Bourguignon, Modern termites inherited the potential of collective construction from their common ancestor. Ecology and Evolution 10, 6775–6784 (2020).

34. A. Buček, et al., Molecular phylogeny reveals the past transoceanic voyages of drywood termites (Isoptera, Kalotermitidae). Molecular Biology and Evolution 39, 1–20 (2022).

35. N. Mizumoto, T. Bourguignon, T. Kanao, Termite nest evolution fostered social parasitism by termitophilous rove beetles. Evolution 76, 1064–1072 (2022).

36. N. Mizumoto, P. M. Bardunias, S. C. Pratt, Complex relationship between tunneling patterns and individual behaviors in termites. The American Naturalist 196, 555–565 (2020).

37. S. F. Light, “Economic significance of the common dry-wood termite” in Termites and Termite Control, C. A. Kofoid, Ed. (University of California Press, 1934), pp. 234–265.

38. O. Kitade, Y. Hayashi, K. Takatsuto, Variation and diversity of symbiotic protist composition in the damp-wood termite Hodotermopsis sjoestedti. Japanese Journal of Protozoology 45, 29–36 (2012).

39. T. Bourguignon, et al., Developmental pathways of Psammotermes hybostoma (Isoptera: Rhinotermitidae): Old pseudergates make up a new sterile caste. PLOS ONE 7, e44527 (2012).

40. P. O. Y. Nkunika, “The biology and ecology of the dampwood termite, Porotermes adamsoni (Froggatt) (Isoptera: Termopsidae) in South Australia,” The University of Adelaide, Adelaide. (1988).

41. G. F. Oster, E. O. Wilson, Caste and Ecology in the Social Insects (Princeton University Press, 1978).

42. D. Cassill, Yoyo-bang: A risk-aversion investment strategy by a perennial insect society. Oecologia 132, 150–158 (2002).

43. L. Bell-Roberts, et al., Larger colony sizes favoured the evolution of more worker castes in ants. Nature Ecology & Evolution 8, 1959–1971 (2024).

44. L. Flintham, J. Field, The evolution of morphological castes under decoupled control. j. evol. Biol. 37, 947–959 (2024).

45. G. J. Thompson, O. Kitade, N. Lo, R. H. Crozier, Phylogenetic evidence for a single, ancestral origin of a “true” worker caste in termites. Journal of Evolutionary Biology 13, 869–881 (2000).

46. T. Bourguignon, R. A. Chisholm, T. A. Evans, The termite worker phenotype evolved as a dispersal strategy for fertile wingless individuals before eusociality. The American Naturalist 187, 372–387 (2016).

47. J. A. L. Watson, J. J. Sewell, “Caste development in Mastotermes and Kalotermes: Which is primitive?” in Caste Differentiation in Social Insects, (Pergamon Press Ltd, 1985), pp. 27–40.

48. F. Legendre, M. F. Whiting, P. Grandcolas, Phylogenetic analyses of termite post-embryonic sequences illuminate caste and developmental pathway evolution. Evolution and Development 15, 146–157 (2013).

49. P. Grandcolas, C. D’Haese, The origin of a ‘true’ worker caste in termites: phylogenetic evidence is not decisive. Journal of Evolutionary Biology 15, 885–888 (2002).

50. L. Revely, P. Eggleton, R. Clement, C. Zhou, T. R. Bishop, The diversity of social complexity in termites. Proceedings of the Royal Society B: Biological Sciences 291, 20232791 (2024).

51. O. Peled, G. Greenbaum, G. Bloch, Diversification of social complexity following a major evolutionary transition in bees. Current Biology 35, 981–993 (2025).

52. J. G. Holland, G. Bloch, The Complexity of Social Complexity: A Quantitative Multidimensional Approach for Studies of Social Organization. The American Naturalist 196, 525–540 (2020).

53. C. Noirot, “The Caste System in Higher Termites” in Caste Differentiation in Social Insects, (Pergamon Press, 1985), pp. 75–86.

54. T. Chouvenc, “A primer to termite biology: Coptotermes colony life cycle, development, and demographics” in Biology and Management of the Formosan Subterranean Termite and Related Species, (CABI, 2023), pp. 40–81.

55. B. L. Thorne, Alate production and sex ratio in colonies of the Neotropical termite Nasutitermes corniger (Isoptera; Termitidae). Oecologia 58, 103–109 (1983).

56. R. Fougeyrollas, et al., Asexual queen succession mediates an accelerated colony life cycle in the termite Silvestritermes minutus. Molecular Ecology 26, 3295–3308 (2017).

57. K.-B. Neoh, C.-Y. Lee, Developmental stages and caste composition of a mature and incipient colony of the drywood termite, Cryptotermes dudleyi (Isoptera: Kalotermitidae). Journal of Economic Entomology 104, 622–628 (2011).

58. P. Luykx, Termite colony dynamics as revealed by the sex- and caste-ratios of whole colonies of Incisitermes schwarzi banks (Isoptera: Kalotermitidae). Insectes Sociaux 33, 221–248 (1986).

59. B. L. Thorne, N. L. Breisch, M. I. Haverty, Longevity of kings and queens and first time of production of fertile progeny in dampwood termite (Isoptera; Termopsidae; Zootermopsis) colonies with different reproductive structures. Journal of Animal Ecology 71, 1030–1041 (2002).

60. J. Korb, S. Schmidinger, Help or disperse? Cooperation in termites influenced by food conditions. Behavioral Ecology and Sociobiology 56, 89–95 (2004).

61. T. Bourguignon, J. Šobotník, R. Hanus, Y. Roisin, Developmental pathways of Glossotermes oculatus (Isoptera, Serritermitidae): At the cross-roads of worker caste evolution in termites. Evolution and Development 11, 659–668 (2009).

62. C. A. Nalepa, Origin of termite eusociality: Trophallaxis integrates the social, nutritional, and microbial environments. Ecological Entomology 40, 323–335 (2015).

63. T. Chouvenc, Eusociality and the transition from biparental to alloparental care in termites. Functional Ecology 36, 3049–3059 (2022).

64. C. A. Nalepa, N. Lo, K. Maekawa, Origin of eusociality in termites: was genetic monogamy essential? Current Opinion in Insect Science 70, 101388 (2025).

65. T. Miura, K. Maekawa, The making of the defensive caste: Physiology, development, and evolution of the soldier differentiation in termites. Evolution and Development 22, 425–437 (2020).

66. J. Korb, K. Schneider, Does kin structure explain the occurrence of workers in a lower termite? Evolutionary Ecology 21, 817–828 (2007).

67. H. Caswell, Matrix Population Models: Construction, Analysis, and Interpretation, Second Edition (Oxford University Press, 2006).

68. I. García-Ruiz, A. Quiñones, M. Taborsky, The evolution of cooperative breeding by direct and indirect fitness effects. Science Advances 8, eabl7853 (2022).

69. I. Stott, S. Townley, D. Carslake, D. J. Hodgson, On reducibility and ergodicity of population projection matrix models. Methods in Ecology and Evolution 1, 242–252 (2010).

70. R Core Team, R: A language and environment for statistical computing. Ver 4.5.2. (2025). Deposited 2025.

71. D. Eddelbuettel, J. J. Balamuta, Extending R with C++: A Brief Introduction to Rcpp. The American Statistician 72, 28–36 (2018).

