## Supplementary material for "Colony phase structure favors permanent worker evolution in juvenile social systems": Table S1 Figure S1-6

This file includes

Table S1
Figure S1-6

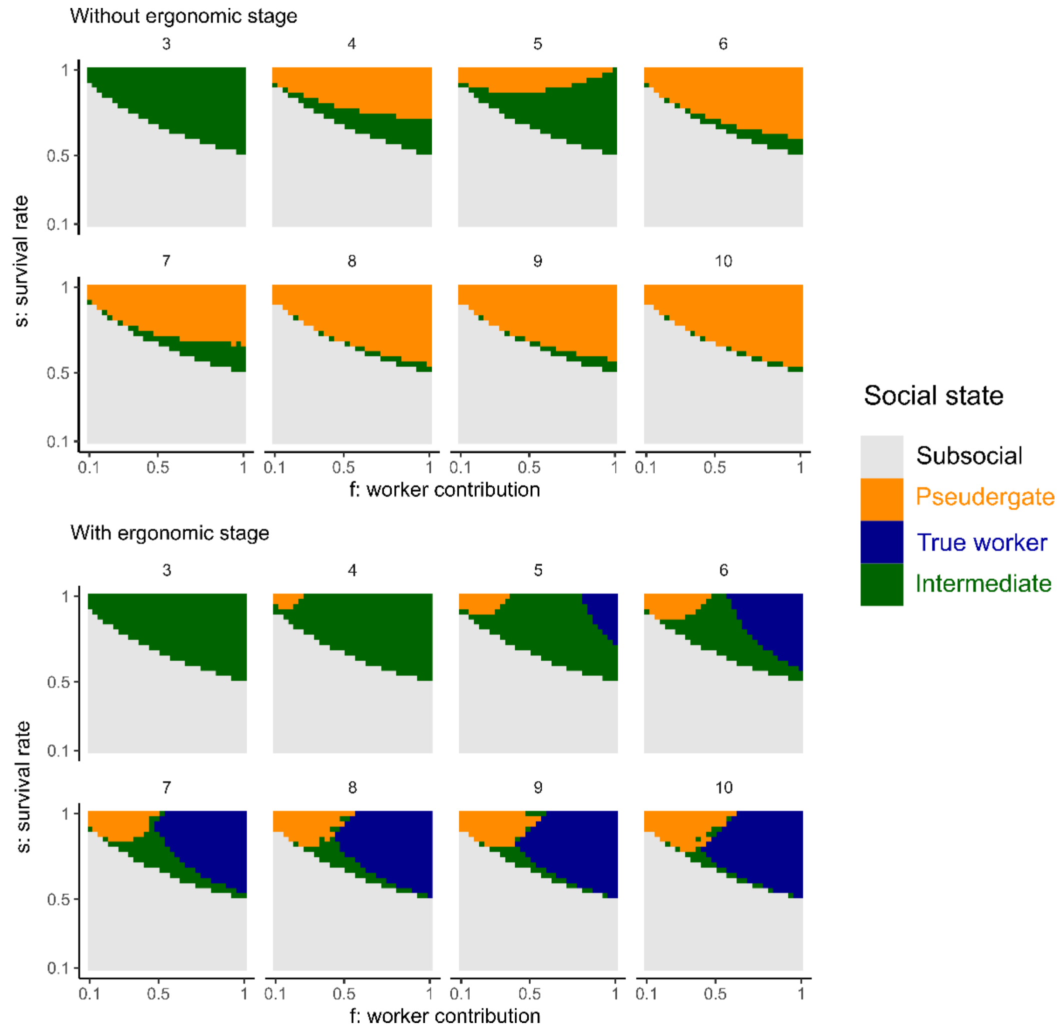


**Figure S1**. Sensitivity analysis of $T_{max}$ in the analytical model with the most advantageous social states. We tested $T_{max}$ in the range of 3-10.

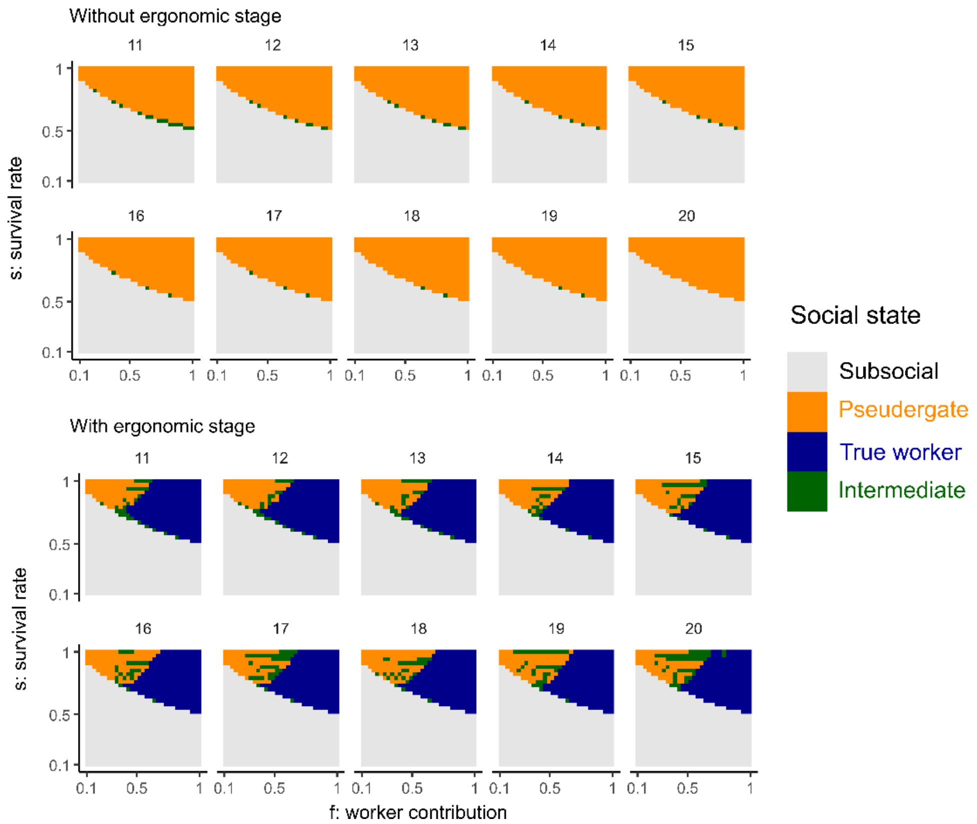


**Figure S2**. Sensitivity analysis of $T_{max}$ in the analytical model with the most advantageous social states, with larger $T_{max}$ values (11-20).

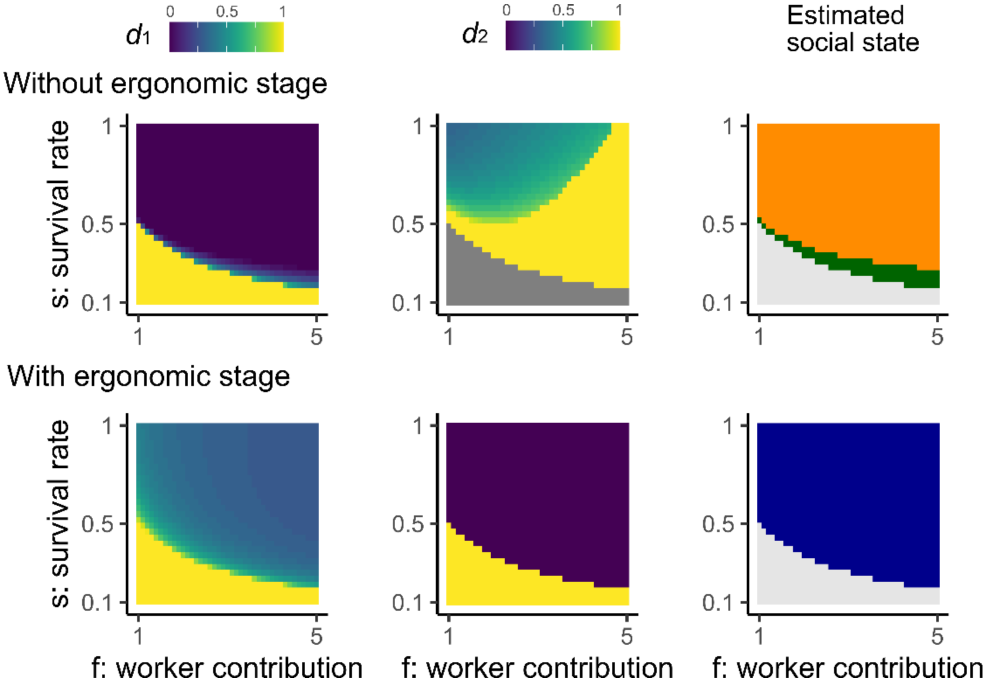


**Figure S3**. Analytical solutions for the most advantageous social strategy (described by $d_1$ and $d_2$) across different parameters, $s$ and $f$, with larger $f$ values.

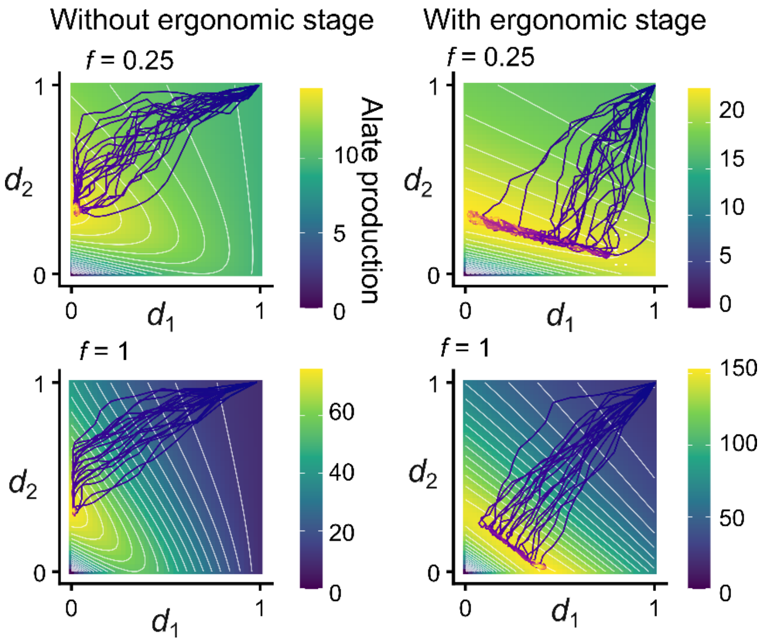


**Figure. S4.** Fitness landscape of social states (represented by $d_1$ and $d_2$) obtained by analytical solutions with parameters for evolutionary simulations ($s$ = 0.95, $T_{max}$ = 10). The evolutionary trajectories correspond to those in Figure 4.

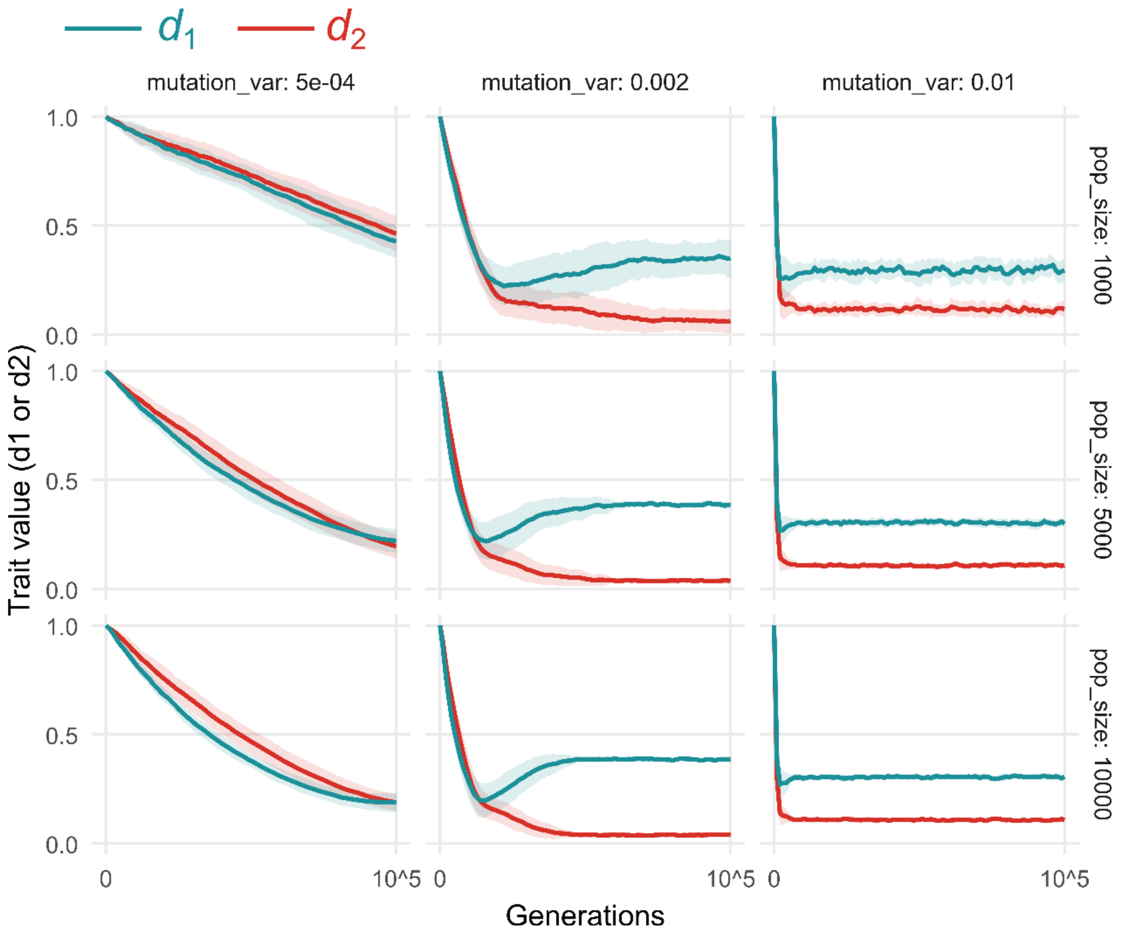


**Figure. S5.** Sensitivity analysis of simulation parameters in evolutionary simulations ($\sigma_{mutation}$ and $N$, population size). The biological parameters are $f$ = 1, $s$ = 0.95, and $T_{max}$ = 10. The results of $N$ = 5,000, $\sigma_{mutation}$ = 0.002 are the same as Figure 4D. Longer generation results for $\sigma_{mutation}$ = 0.0005 are in Figure S6.

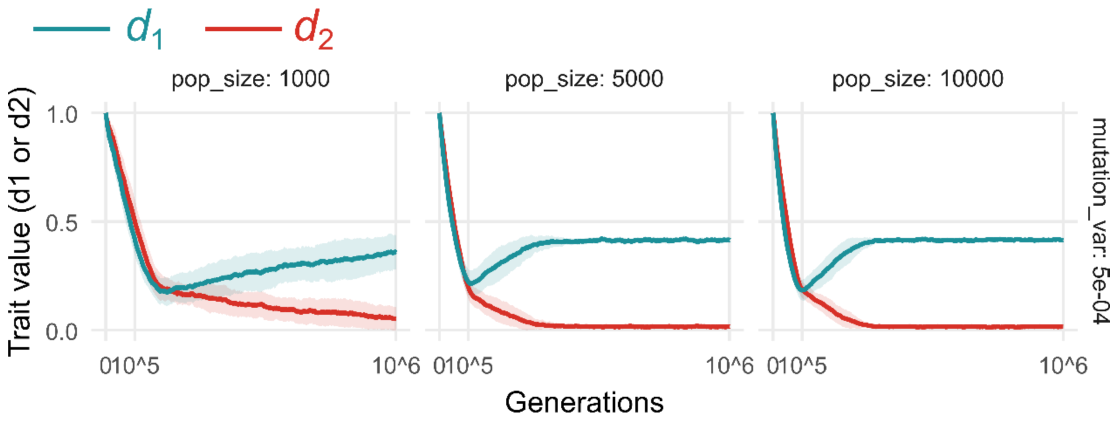


**Figure. S6.** Longer trajectories for the evolutionary simulations with $\sigma_{mutation}$ = 0.0005.

**Table S1**. Parameters and variables

| | | |
|---|---|---|
| $d_1$ | [0,1] | Probability for brood to become alate |
| $d_2$ | [0,1] | Probability for worker to become alate |
| $s$ | [0,1] | Survival rate within the nest |
| $f$ | [0,5] | Worker contribution to brood production |
| $T_{max}$ | [1,10] | Colony life span |
